# Down regulation of *Engase* in *Caenorhabditis elegans* may improve its stresses adaptivity

**DOI:** 10.1101/2024.07.01.601486

**Authors:** Xinrong Lu, Yongliang Tong, Mengting Wu, Shaoxian Lyu, Jiale Fan, Junyu Zheng, Lin Zou, Danfeng Shen, Lin Rao, Linlin Hou, Cuiying Chen, Xunjia Cheng, Guiqin Sun, Zhiyong Shao, Li Chen

**Author notes:** Correspondence auther: Li Chen, Zhiyong Shao, Guiqin Sun. These authors contributed equally to this work (Xinrong Lu, Yongliang Tong).

## Abstract

Endo-beta-N-acetylglucosaminidase (ENGASE) is one of the key enzymes involved in the regulation of structure and function of glycoproteins. It is conserved from prokaryotic to eukaryotic cells. Although their activities *in vitro* and applications have been well studied, the biological function of ENGASE remains to be illustrated. In this study, we analyzed the molecular and physiological function of *Engase* from *Caenorhabditis elegans* homolog *eng-1*(*CeEngase*). We found that *CeEngase* knockout or knockdown increased the environmental stresses adaptability, such as heat stress and osmotic stress. Preliminary glycomics analysis showed that the basement membrane proteins of extracellular matrix may be the main targets of CeENGASE. In addition, CeENGASE may selectively prefer to N2H7 glycans on glycoproteins. In conclusion, our data illustrated that the defection and/or down regulation of *CeEngase* may provide a beneficially adaptation for stresses.

## 1. Introduction

Protein N-glycosylation is a dynamic post-translational modification process found in eukaryotic cells. It is involved in proteins’ folding trafficking and digestion, cell surface signaling, microbial-host interactions and immune responses, [1, 2]. Its alterations may reflect physiological and/or pathological condition and may serve as potential therapeutic targets [3].

ENGASE is an endo-deglycosylase for N-glycans on glycoproteins and free oligosaccharides (fOSs). It hydrolyzes a β-1,4 glycosidic bond between two N-Acetylglucosamines (GlcNAc) in core pentasaccharide [4, 5]. In addition to its hydrolysis activity, a reverse synthesis activity was also reported and has been used in engineering for glycoproteins with uniformed N-glycan [6, 7].

Previous studies reported that *Engase* was a potential theraptic target for *Ngly1* deficient diseases, which often result in embryonic lethality and can be rescued by *Engase* knockout [3, 8]. In shrimp, the cumulative mortality and viral copy number were significantly decreased when *PmEngase* (*Penaeus monodon Engase*) was silenced [9]. *Trichoderma atroviride TaEngase18B* (*Trichoderma atroviride Engase*) knockout strain grew faster and was more resilient to abiotic stresses under stress conditions for cell wall, such as 1mol/L NaCl or 0.025% SDS [10]. Accumulated evidences suggest that *Engase* inhibition might play a protective role in stress conditions.

*Caenorhabditis elegans* is an ideal multicellular eukaryotic animal model for complete life cycle and whole single cell analysis. Its genetic and developmental background are well characterized and the analytic systems for molecular analysis at single cell level are well established. In addition, it shares many basic biological processes, such as aging, digestion, and stress adaptations with human beings [11]. *CeEngase* (*Caenorhabditis elegans eng-1*) was discovered and characterized by Kenji Yamamoto *et al* based on its sequence homology to *Endo-M* from *Mucor hiemalis* [12]. Following this study, *CeEngase* had been cloned, prepared and subjected to in vitro assays for its enzymatic activity, substrate specificity and optimal reaction conditions. *CeEngase* deletion mutant was constructed by using conventional mutagenesis. In addition, fOSs and some basic biological indicators were examined under normal conditions [13, 14].

In this study, a frameshift mutant in exon 2 of *CeEngase* was constructed and subjected to biological and biochemical analysis. The survival rates at different stress conditions of the mutant were measured and compared to wild type. The total proteins were collected and subjected to a glycomic assay and the results of the glycomic assay were analyzed and presented.

## 2. Materials and methods

### 2.1 Strains and maintenance

All *C. elegans* strains were cultured on nematode growth media (NGM) plates with OP50 standard food [15]. All assays were performed at 21°C unless otherwise stated. Worms were used in this study at day 1 adult stage unless specified. Detailed information for strains was listed in Supporting Information: Table S1.

### 2.2 *CeEngase* gene (*eng-1*) knockout

CRISPR-Cas9-guided genome editing was performed as previously described [16]. sgRNA for *CeEngase* gene (*eng-1*) knockout (5’-CACTCGAAGAGCTCTGGAGT-3’) was designed by using benchling. The sgRNA was inserted into pDD162 vector [16] for the plasmid (pLXR1). Successful knockouts were screened by PCR and sanger sequencing.

Genomic DNA was sequenced (primer: 5’-GCGGGATCTGGAAATTGTGCTATAT-3’) for knockout. A second around single/sequencing was performed for diploid knockout. Sequencing confirmed line was outcrossed 4 times with wild-type (N2) worms and the confirmed knockout after outcross was named as *LXR1*. Detailed information for plasmids and primers was listed in Supporting Information: Table S2.

### 2.3 Confirming knockout at protein expression level

Using genomic DNA as template, a fragment containing the promotor (2000bp) and coding sequence (1302 bp) of *CeEngase* was amplified, and inserted into pSM vector [17]. After confirming the construction by sequencing, GFP was inserted at upstream of *CeEngase* stop codon. In this plasmid (pLXR2) the expression of CeEngase/GFP (*PCeEngase*::CeEngase::GFP) was similar as the one on chromosome and its expression could be monitored by GFP’s fluorescence. Site directed mutagenesis was conducted to generate the 4 base deletion detected on *PCeEngase*::CeEngase::GFP (pLXR3). Detailed information for plasmid and primers was listed in Table S2.

### 2.4 Fluorescence and confocal microscope imaging

Worms with GFP expression were anesthetized with 50 mM Muscimol on 3% agarose pads on slides. Images were captured with Andor Dragonfly Spinning Disc Confocal Microscope with 10× objective lens, with 488 nm laser. Image processing was performed with Imaris 10.0/Image J software.

### 2.5 Lifespan assay

Lifespan assays were performed following a standard protocol [18]. Briefly, synchronized larval stage 4 (L4) worms were placed on fresh NGM plates and incubated. Survived worms were counted daily. Those worms that did not respond to external mechanical stimuli were counted as dead. Bagged, internal hatching or missing worms were counted as censored. Worms were transferred to new NGM plates every other day. 60 worms were checked for each test. Three tests were performed for this lifespan assay. Kaplan-Meier survival analysis with log-rank test were used to analyze the lifespan of the worms.

### 2.6 Motoricity assay

The assay was used to check the impact of a specific treatment on worm’s mobility by measure the head-swing ability in liquid. The reported protocol was followed [19]. Briefly, synchronized worms were placed on fresh NGM plates. 200 μL of M9 solution was dripped into the plates to help worms swing freely. The times of head swings in 10 seconds were counted. 20 worms were checked for each test. The tests were repeated three times.

### 2.7 Osmotic stress resistance assay

Osmotic stress resistance assays were performed following a published protocol with slight modifications [20]. Synchronized L4 stage worms were placed on fresh NGM plates for 24 h. Day 1 adults were transferred from standard 50 mM NaCl NGM plates to high salt NGM plates (≥400 mM NaCl) and incubated at 21°C for 24 h. Worms that did not respond to external mechanical stimuli were counted as unsurvived and sensitive to osmotic stress. 60 worms were checked for each test, and the test was repeated three times.

### 2.8 Heat stress assay

Heat stress assays were performed following the instruction that was slightly modified [21]. Synchronized L4 stage worms placed on fresh NGM plates were incubated at 21°C for 24 h. Day 1 adults were subjected to high temperature stress at 35°C for 3 h or 7 h, respectively. Those worms that did not respond to external mechanical stimuli after the treatment were counted as sensitive. The percentage of survived worms were scored. 60 worms were checked for each test, and the test was repeated.

### 2.9 RNA interference

RNAi feeding bacteria and plates were prepared following the previous study [22]. Eggs were collected from 20 synchronized day 1 adults for 2 h and fed them with RNAi bacteria. Day 1 adults were incubated at 35°C, 11 h for stress assay. The survival rates were measured.

### 2.10 Structure prediction and alignment

The structures of CeENGASE (Q8TA65) and hENGASE (Q8NFI3) were predicted by AlphaFold 3 respectively [23]. The alignment of the two was obtained with open accessed ChimeraX. The display option in Chimera was used to mark critical residues.

### 2.11 Protein expression and preparation

*CeEngase* (*eng-1*) gene was cloned into pET28a vector (Genscript). The constructed pET28a-*eng-1* plasmid was transformed into *E. coli* BL21 (DE3) (Tiangen). Single clones then were cultured in Luria-Bertani medium, and the expression induced of proteins by IPTG (1mM) at 21°C for 16 h. The proteins were purified using Ni Sepharose ^TM^ 6 Fast Flow (GE Healthcare).

### 2.12 Enzyme activity against glycoprotein

This assay was performed following a published protocol with slight modifications [24]. In brief, a 200 μL reaction system in 50 mM sodium acetate rection buffer (pH 5.5) contain 100 μg RNaseB (Sigma), 100 μg CeENGASE and 150 μM Rabeprazol sodium (Topscience) was incubated at 21°C for 24 h. The reaction was terminated at 100°C for 10 min. After a spin (10,000 g 10 min), the free glycans in supernatant generated in the reaction were collected, purified by 3 kD ultrafiltration, processed to remove salt by Supelclean^TM^ EVNI-Carb^TM^ SPE Tube (Sigmaaldrich), condensed by lyophilization, dissolved in ddH2O, and subjected to MALDI-TOF MS analysis.

### 2.13 Rabeprazole feeding assay

The assay was performed following the protocol with minor modifications [25, 26]. Synchronized L4 stage worms were treated with 1 mM Rabeprazole sodium [27] and solvent control (DMSO) in OP50 food, respectively. After one and half day’s treatment, plates containing these adults were incubated at 35°C, 7 h for stress assay. The survival rates were measured.

### 2.14 Detection and analysis of glycopeptides in *C. elegans*

Around 5000 day 1 adult worms on NGM plates were collected, suspended in M9 buffer, and washed three times with the same buffer. The precipitated worms in M9 were collected, treated in RIPA lysis buffer (Beyotime) plus PMSF (100 mM, 1:1000) (Beyotime) on ice for 3×5 min with vortex in interval. The sample were further processed by ultrasonication (10×5 seconds) Ultra-high centrifugal speed (13,000 g, 10 min), supernatant collection, reductive alkylation, and acetone precipitation [28].

The precipitates were solubilized in 25 mM ammonium bicarbonate solution, trypsinized at an enzyme/protein ratio of 1/50, and digested at 37°C overnight. The digested peptides were first purified by C18 desalting column (Waters) and then processed for N-glycopeptide enrichment using the ZIC-HILIC Glycopeptide Enrichment Kit (Mercks). The enriched N-glycopeptides were lyophilizated, solubilized in 0.1% formic acid solution, and analyzed by Thermo Fisher Easy-nano LC II system. Mass spectrometry raw data were analyzed using proteome discoverer software (Thermo Fisher) and pGlyco3.0 software [29]. Quantitative analysis was performed by peak finding, extraction and intensity quantitation functions in pGlycoQuant [30].

Wild type (N2) and *LXR1* worms were subjected to this assay, and three technical replicates were performed for each set of samples.

### 2.15 Statistical analysis

All assays were statistically analyzed using GraphPad Prism 9.5.0. Comparison of data between the two groups was analyzed using an unpaired two-tailed Student’s t test. Values are expressed as mean ± standard error of the mean.

## 3. Results

### 3.1 Construction and validation of a *CeEngase* knockout strain

Briefly, pLXR1 constructed for knockout was injected at gonadal cell together with *Pmyo-3*::*mCherry* and P*myo-2::mCherry*. The genomic DNA of offsprings were made and subjected to a PCR reaction covering the first three exons of *CeEngase*. The PCR products were sequenced for candidate mutants. After 4 rounds of screening a knockout mutation with a 4 base deletion inside the second exon of *CeEngase* was selected and named as *LXR1* for further studies (Fig 1A). A fast PCR reaction that was positive only for wild type genome and negative for *LXR1* was designed for the quality control of *LXR1* (Fig S1A). The impact of the frameshift mutation in *LXR1* were further confirmed at the mRNA level by a transcriptome sequencing analysis (Fig S1B) and at protein level by an in vivo *PCeEngase*::CeEngase::GFP assay (Fig 1B).

**Figure 1.**
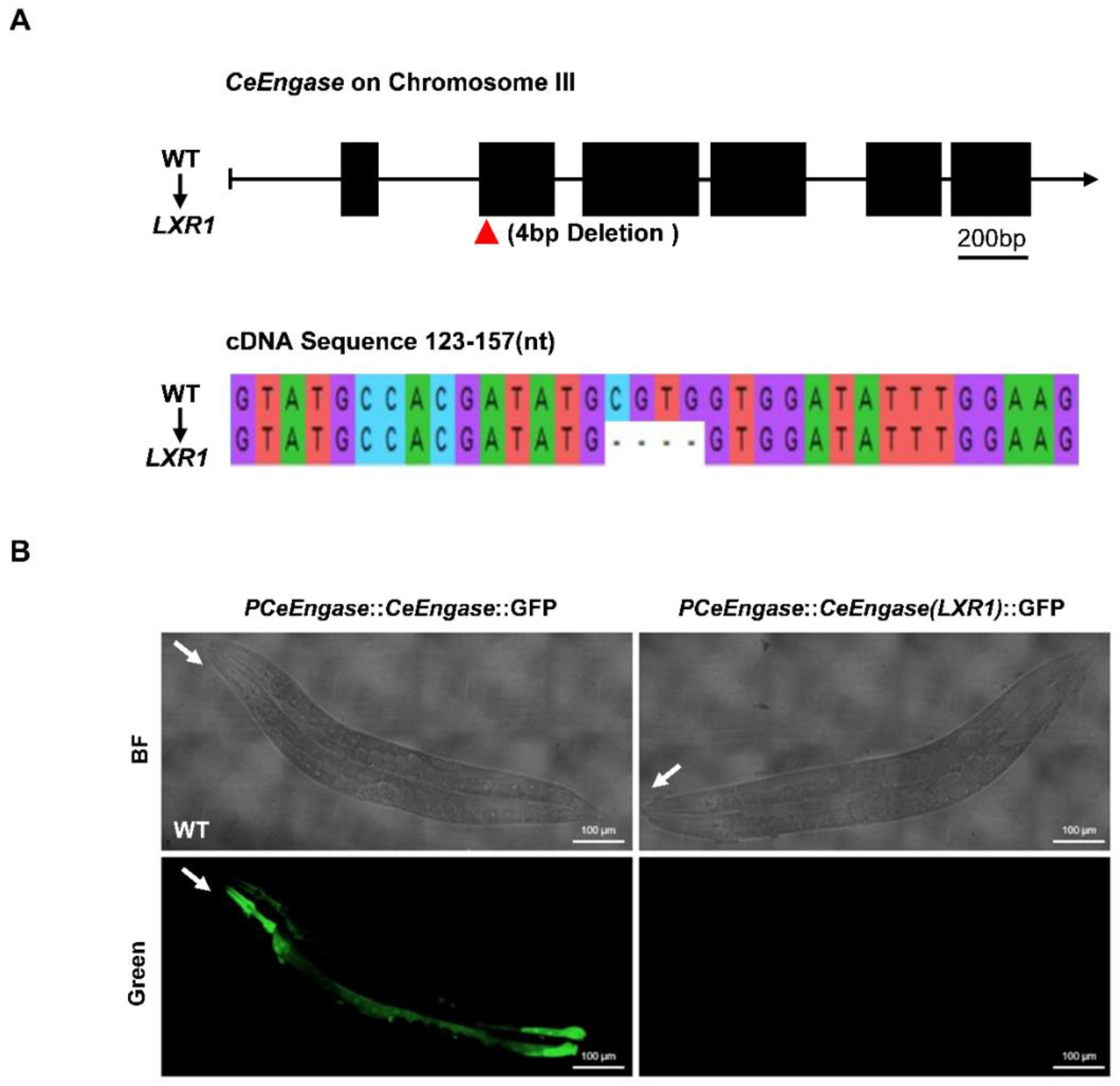
Construction and validation of a *CeEngase* knockout strain. (A) Schematic drawing of *CeEngase* gene in wild type and *LXR1* allele. Exons are depicted as filled black boxes. The genomic deletion in *LXR1* allele was indicated by arrow. (B) Frameshift mutation in *LXR1* was confirmed by an GFP fusion expression assay. The two plasmids (pLXR2; pLXR3) were injected into the wild type worm for GFP expression respectively. The heads of the worms were indicated with white arrow.

### 3.2 Knockout of *CeEngase* improved stress resistance of worms

To explore the impacts of *CeEngase* on *C. elegans*, the lifespan and motricity were measured. Comparing to the wild type, the lifespan of *LXR1* was improved. The median lifespan of *LXR1* was extended 2 more days than the wild type (*p*<0.05) (Fig 2A). In addition, the body movement of worms were measured as an indicator for healthy living. Head swings of worms in M9 solution were counted. A higher frequency was observed in *LXR1* (*p*<0.01) at adult day 5 (Fig 2B). Together these results show that *CeEngase* knockout has a positive effect on worms.

**Figure 2.**
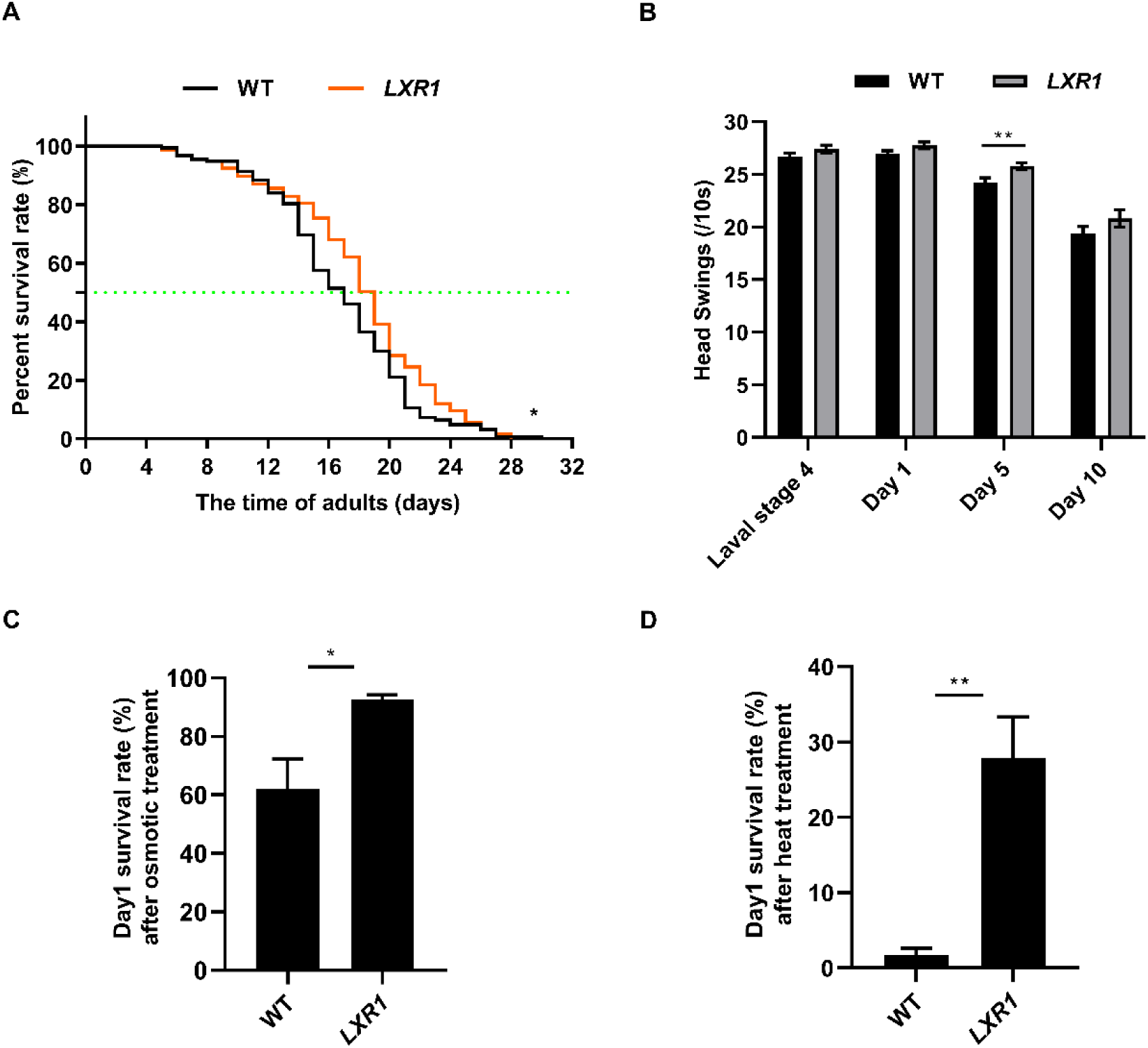
Knockout of *CeEngase* improved stress resistance of worms. (A) The survival curves of worms. (B) The frequencies of head swing in M9 solution at larval stage 4, day 1, day5, and day 10. (C) The survival rate of *C. elegans* under osmotic stress. (D) The survival rate of *C. elegans* after heat treatment at 35°C for 7 h.

To address the effect of *CeEngase* knockout to stress, *LXR1* and wild type were subjected to high osmotic and temperature conditions respectively. First, we tried high osmotic conditions by incubating day 1 adults on NGM plates containing high salt (≥400 mM NaCl). After incubated at 21°C for 24 h, the survival rates were calculated. The results showed that under osmotic stress, the mean survival rate of *LXR1* was 30.6% more than wild type (*p*<0.05) (Fig 2C). Next, to investigate whether *CeEngase* knockout enhanced worms’ resistance to heat. Day 1 adults were incubated at 35°C for 7 h. The results showed that the mean survival rate of *LXR1* treated at 35°C for 7 h was 26.2% more than wild type (*p*<0.01) (Fig 2D). Both of these experiments suggest that knockout of *CeEngase* increases worms adaptivity to stress.

### 3.3. Knockdown of *CeEngase* improved heat stress resistance of worms

To further validate whether the down regulation of *CeEngase* had similar stress tolerance phenotype, *CeEngase* RNAi was performed. Firstly, the knockdown efficiency of *CeEngase* RNAi was confirmed (Fig 3A). Then, the stress assay was conducted at 35°C for 11 h. We found that the survival rate of *CeEngase* RNAi worms was significantly increased (25.3%, *p*<0.01) (Fig 3B). This result implied that reducing the expression of *CeEngase* could improve the resistance of *C. elegans*. The result may also be concluded that the phenotypes obtained in *LXR1* worms were mainly through the loss of *CeEngase* but not the other background issues.

**Figure 3.**
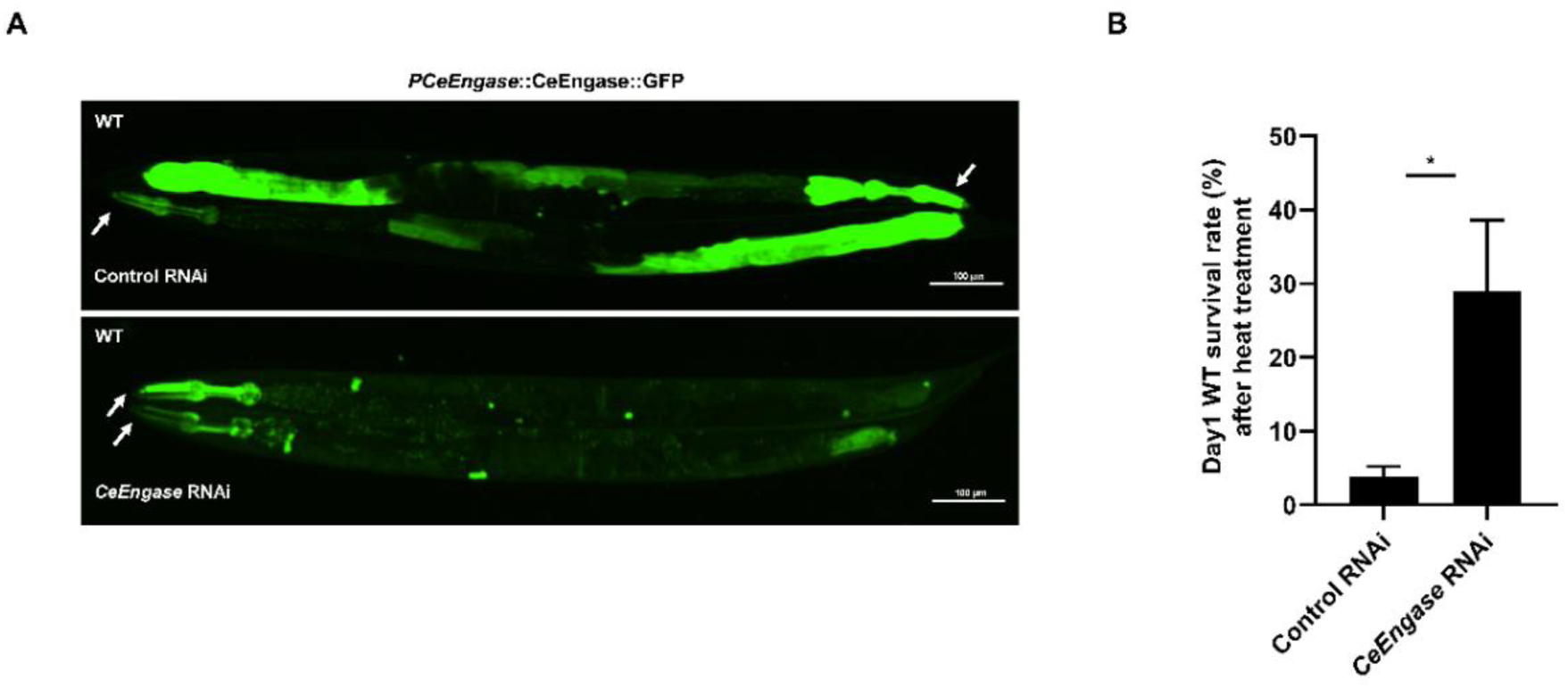
Knockdown of *CeEngase* improved the heat stress resistance of worms. (A) The expression of *PCeEngase*::CeEngase::GFP was down regulated by *CeEngase* RNAi. (B) Knockdown of *CeEngase* increased the survival rate of *C. elegans* under stress.

### 3.4 Structure of CeENGASE was predicted and aligned with hENGASE

For structure flexibility of its substrate and other reasons, the crystal and/or freeze electronic microscope structure of ENGASE remains elusive. The CeENGASE and hENGASE structures were predicted by AlphaFold 3 (Fig 4A; 4B) and aligned by ChimeraX. A common domain for substrate binding and enzymatic hydrolase were presented (Fig 4C). His181 and Asn235 (purple) two conserved residues in the catalytic pocket of hENGASE were identified by mutation analysis (data not shown). The matched residuals in CeENGASE are His99 and Asn152 respectively (beige) (Fig 4C).

**Figure 4.**
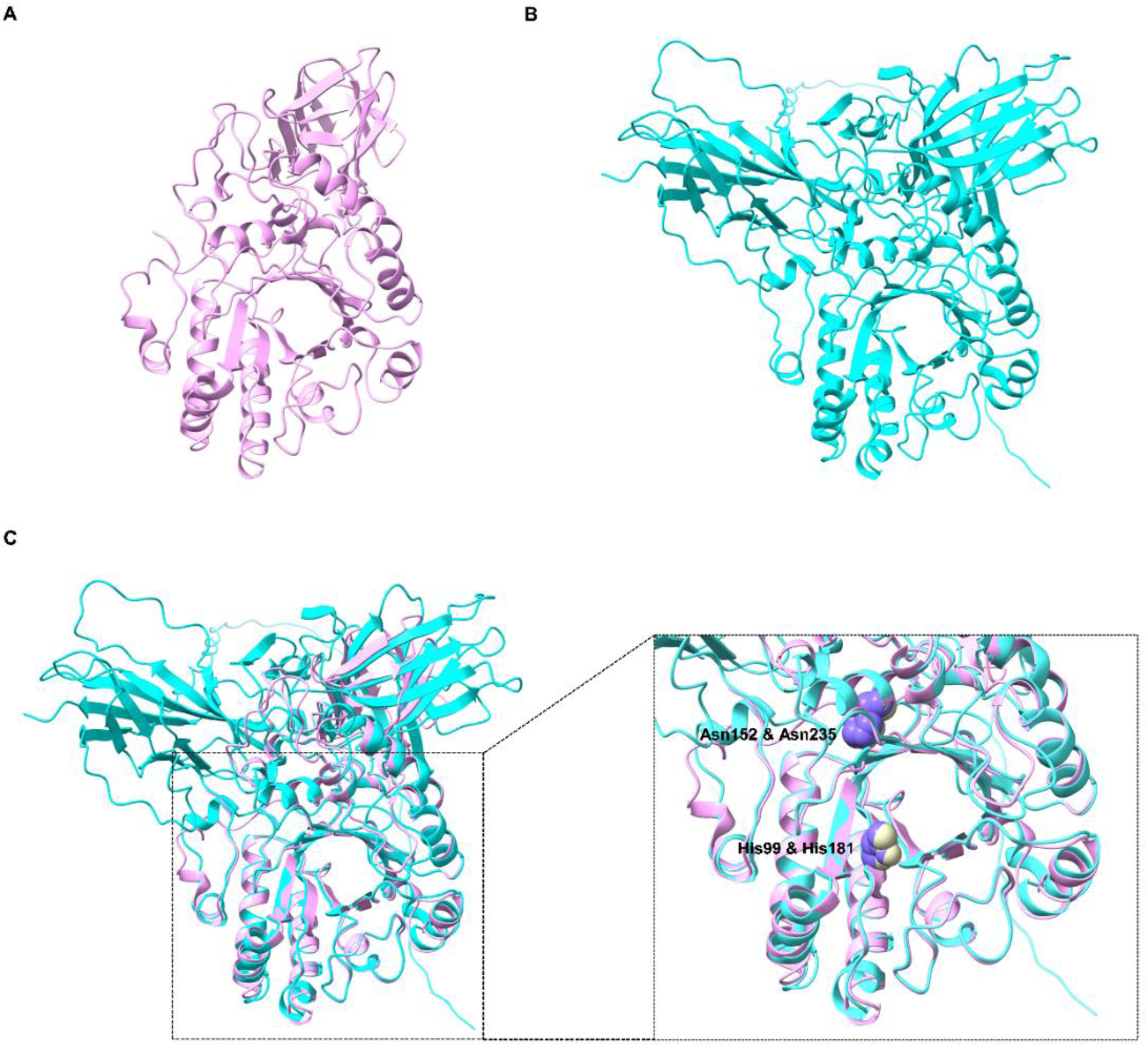
Structures of CeENGASE and hENGASE. (A) Structure of CeENGASE. (B) Structure of hENGASE. (C) Structures alignment of CeENGASE and hENGASE. Two critical residuals in the hydrolase domain of CeENGASE and hENGASE were marked.

### 3.5 Rabeprazole inhibited CeENGASE *in vitro* and improved worms stress adaptivity *in vivo*

Early studies reported that hENGASE was inhibited by Rabeprazole *in vitro* [27]. Given the structure and function similarity between hENGASE and CeENGASE, we wanted to know if Rabeprazole can inhibit CeENGASE. We first tested the role of Rabeprazole in inhibiting CeENGASE *in vitro*. CeENGASE expressed in prokaryotic system was prepared and subjected to enzyme activity assay. RNaseB was used as the substrate, and N1H7 (GlcNAc1Hexose7), one of the products generated by CeENGASE was detected (Fig 5A). The activity was significantly blocked by Rabeprazole (150 μM) (Fig 5A), suggesting that Rabeprazole also inhibited the enzymatic activity of CeENGASE.

**Figure 5.**
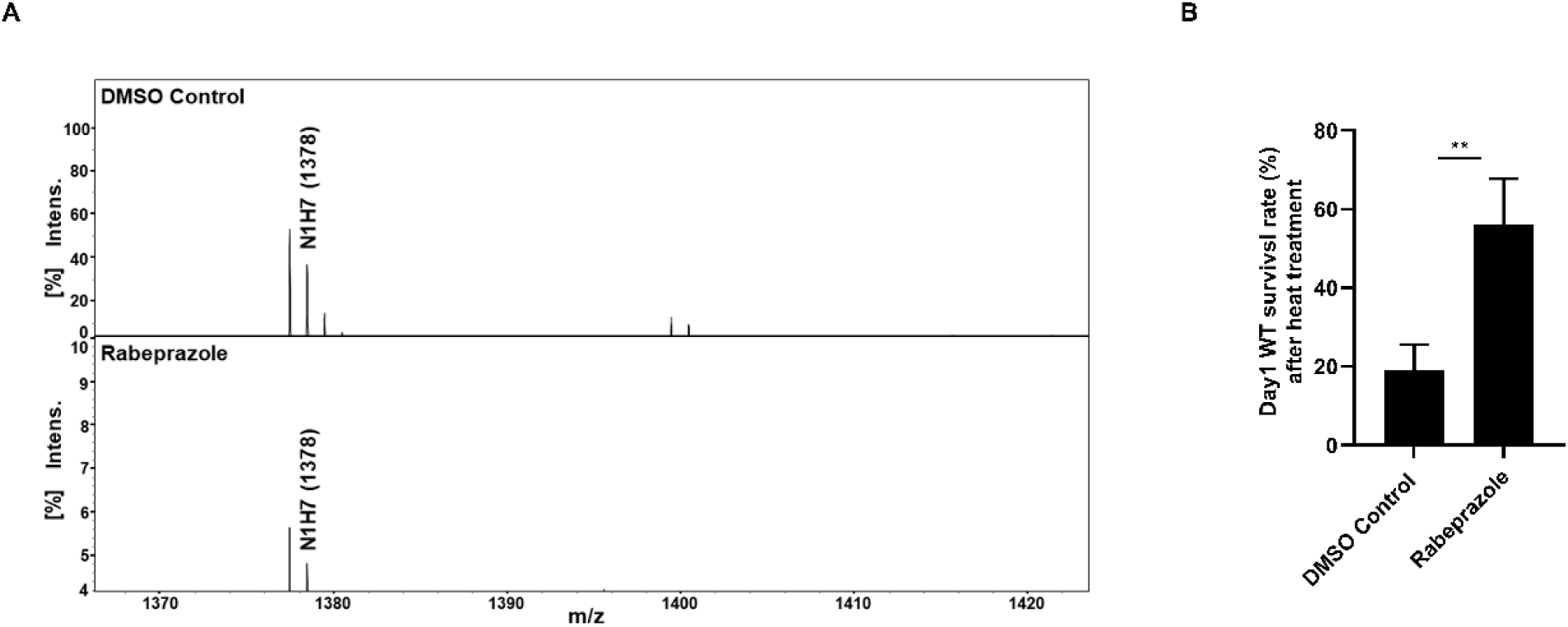
Rabeprazole inhibited CeENGASE *in vitro* and improved worms stress adaptivity *in vivo*. (A) *In vitro* activity of CeENGASE was inhibited by Rabeprazole. (B) The survival rate of *C. elegans* under stress was improved by Rabeprazole.

Next, we wanted to test the role of Rabeprazole in inhibiting CeENGASE *in vivo*. To address this question, we analyzed the stress resistance phenotype of worms treated with Rabeprazole. Stress assay was performed at 35°C for 7 h to check the effect of Rabeprazole treatment. The result showed that the survival rate was significantly increased (36.9%, *p* < 0.01) in Rabeprazole group (Fig 5B). This result indicated that Rabeprazole inhibited CeENGASE function and further support the hypothesis that elimination or inhibition of CeENGASE function increased the resistance of *C. elegans* to stress.

### 3.6 A comparative glycomics analysis for N-glycans from *LXR1* and wild type (N2)

A glycomics analysis was conducted to identify the targets of CeENGASE in worms. Proteins from wild type and *LXR1* were prepared, treated with trypsin and subjected to LC-MS. The data was analyzed by pGlyco3.0 and pGlycoQuant for glycopeptides. A total of 672 and 664 N-glycopeptides were detected in wild type and *LXR1* respectively. Among which 11 were found only in wild type and 3 were found only in *LXR1* (Table 1). For the glycopeptides found in both groups, the ratio of *LXR1*/WT were calculated. The ones significantly high (≥7) and low (≤ 0.05) were listed (Table 1). We noticed that in the Table 1 glycopeptides generated from glycoproteins associated with basement membrane (BM) were significantly enriched (22.1% up and 21.90% down), among which LAM-1, LAM-2, EPI-1, PAT-2, PAT-3, and MIG-6 were included (Fig 6). This result indicated that BM proteins were sensitive to the down-regulation of CeENGASE.

**Table 1.**
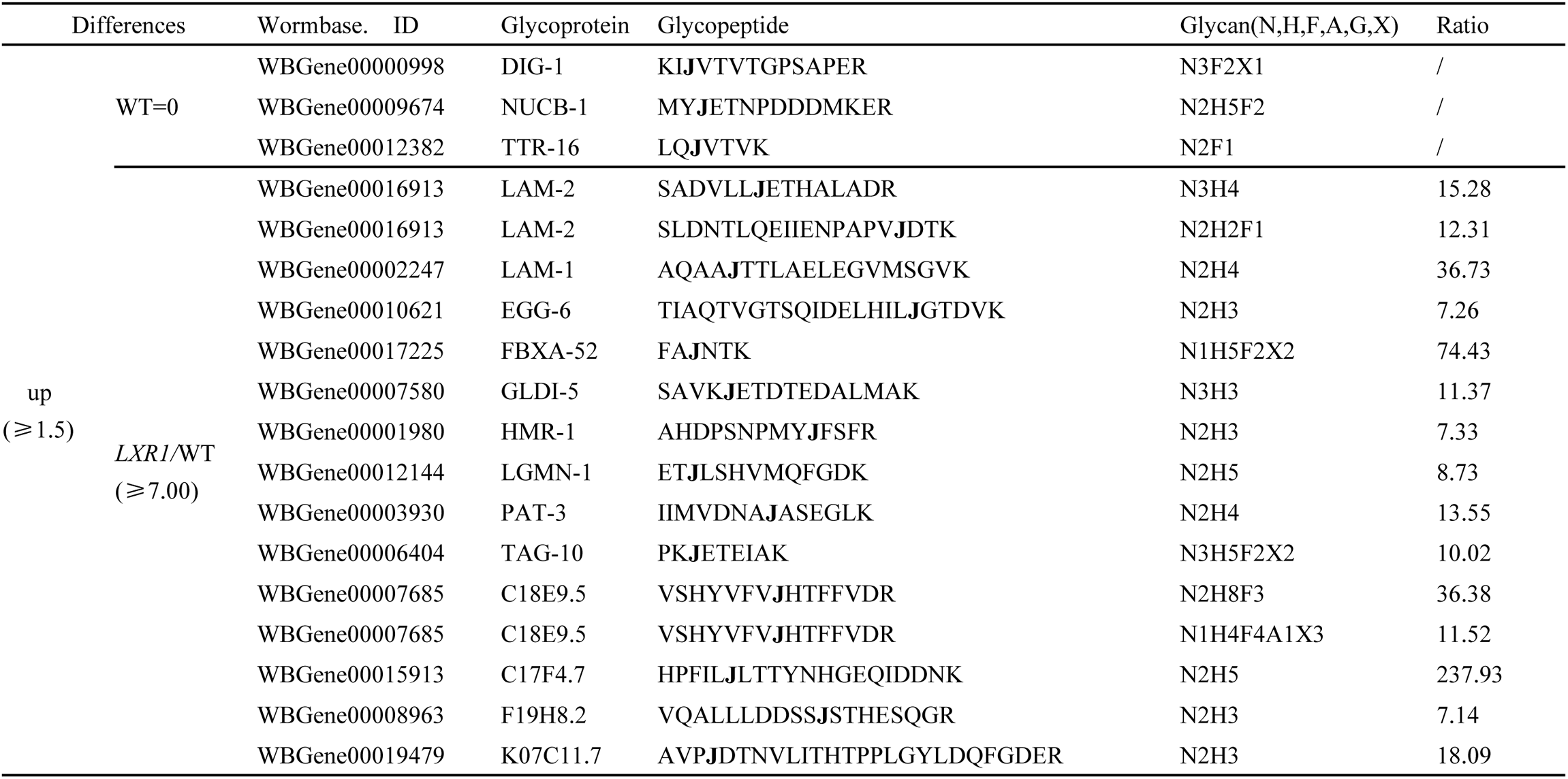
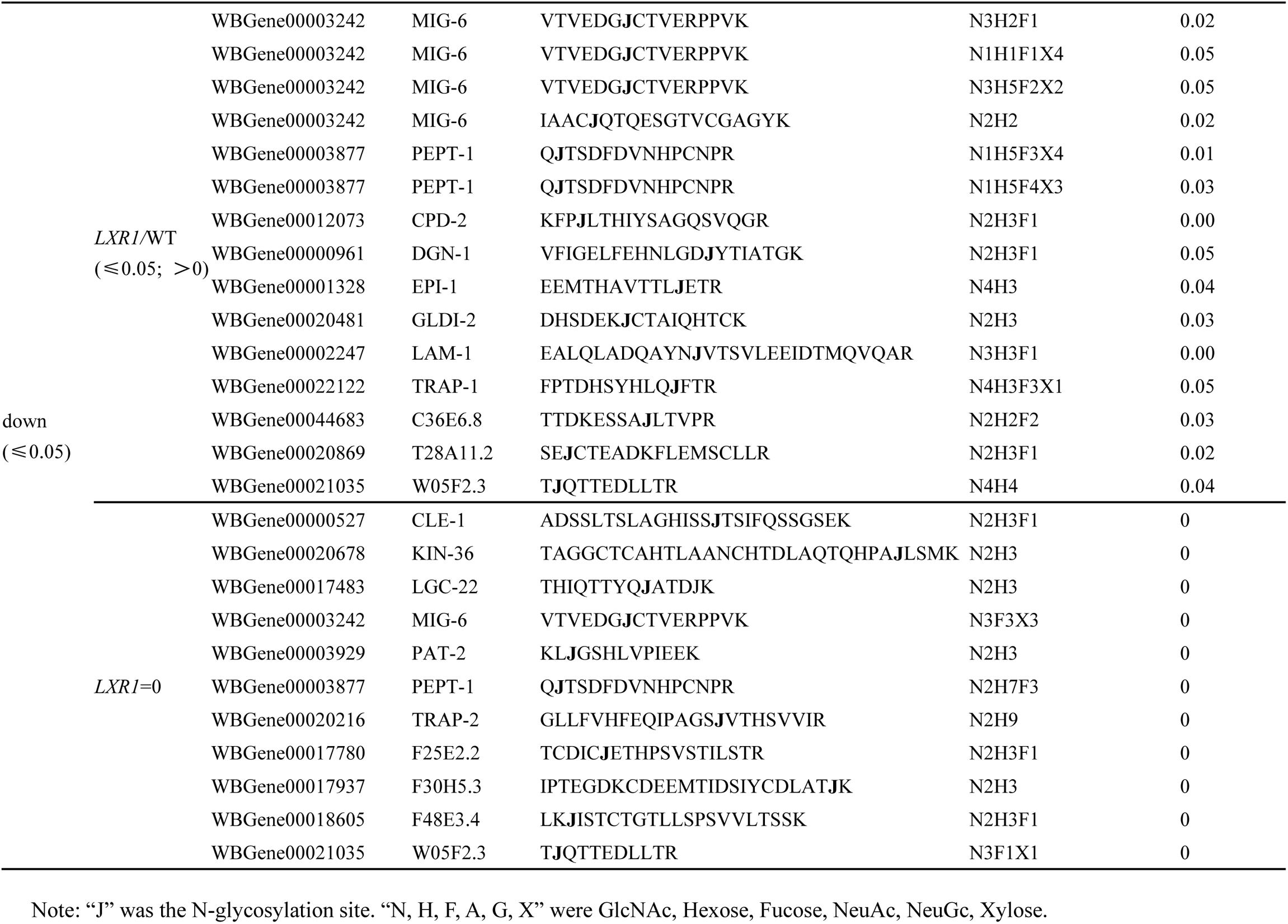
The N-glycopeptides which were significantly changed in *LXR1*.

**Figure 6.**
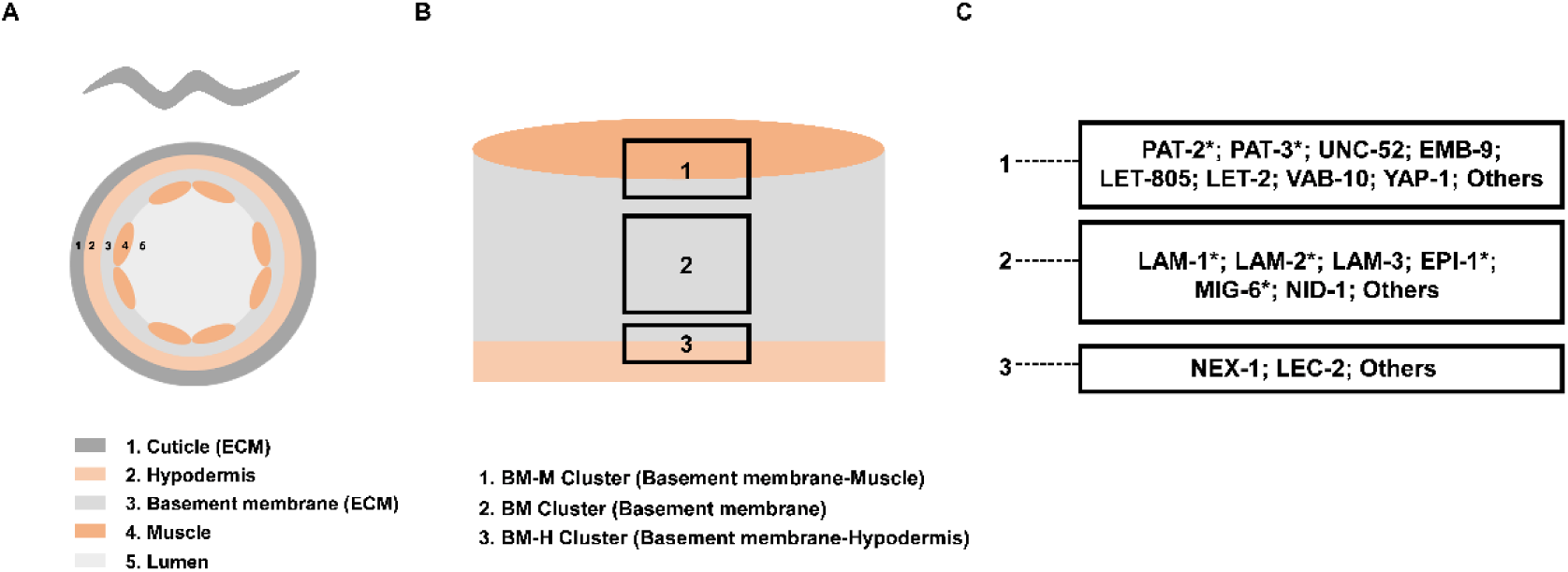
Detected glycopeptides in basement membrane and their sensitivity to CeENGASE regulation. (A) Cross-section of worms. (B) Basement membrane proteins could be divided into three clusters. (C) Reported Proteins in each cluster and the ones containing significantly changed glycopeptides in *LXR1*. The significant changes in glycoproteins were marked by “*”.

### 3.7 Impact of *CeEngase* knockout on profile of detected N2HX glycopeptides

The glycopeptides detected from wild type and *LXR1* worms in glycomics assay were subgrouped by N2HX and compared. In the 675 (672+3, 664+11) detected glycopeptides, 274 were in the form of N2HX. These N2HX were further divided into 10 sungroups. In each subgroup, the percentile of significantly up- and down-regulated by *CeEngase* was calculated and listed (Table 2). The average percentile of up and down for all subgroups was 22% and 19% respectively. Compared to the average, N2Hl was significantly higher (40.00%) in the up group and N2H7 was significantly lower (9.09%) in down group. The amplitude of variation in each subgroup was defined by % of detected ups divided by % of downs. The result of this calculation indicated that N2H7 was more sensitive to CeENGASE regulation. This prediction was consistent with the *in vitro* results generated in this study (Fig 5A) and early reports [12, 13].

**Table 2.** Changes of N2HX glycopeptides in *LXR1*.

| glycopeptides | up ($\geq 1.5$) | down ($\leq 0.5$) | up/down |
| --- | --- | --- | --- |
| N2H0 | 20.00% | 0.00% | / |
| N2H1 | 40.00% | 20.00% | 2.00 |
| N2H2 | 26.67% | 20.00% | 1.33 |
| N2H3 | 18.79% | 24.24% | 0.78 |
| N2H4 | 28.57% | 21.43% | 1.33 |
| N2H5 | 21.74% | 17.39% | 1.25 |
| N2H6 | 11.11% | 22.22% | 0.50 |
| **N2H7** | **27.27%** | **9.09%** | **3.00** |
| N2H8 | 0.00% | 25.00% | 0.00 |
| N2H9 | 26.32% | 21.05% | 1.25 |

## 4. Discussion

Protein N-glycosylation was a comprehensive post translational process accomplished by a series of sequential reactions, including one catalyzed by ENGASE. Early studies indicated that down regulation of *Engase* was a cellular response to biological stresses, such as *Ngly1* defect [3, 8, 31] and viral infection [9]. Known as one of the key soil organisms [32], *C. elegans* had being exposed to frequent heat, osmotic and other environmental stresses. Early studies indicated that heat shock proteins (HSP) are major factors responsible for heat adaptations via rescue damaged proteins for protein homeostasis [33], and tens of factors on extra extracellular matrix such as laminins are critical for its survival [34]. In addition, more than 70 genes, including *daf-2, daf-16, sir-2.1,* and others are implicated in longevity regulation [35]. In this study, we demonstrated that a mild but significant improvement of all these three health indexes was achieved by down-regulation of *CeEngase*. We concluded that *CeEngase* down-regulation may represent a general stress management strategy. As the biological quick recognition information carried by N-glycans on glycoproteins, such as signals for folding trafficking and localization were removed by ENGASE, our observations might suggest that ENGASE was a resource for protein toxicity. More studies were required to explore the negative protein toxicity of ENGASE and its positive biological impact that remained elusive.

A *CeEngase* knockout with a 4 base frameshift deletion in the first exon was generated and confirmed at DNA, RNA, and protein levels (Fig 1). In addition, the deletion was independently confirmed by the data generated from a transcriptomic analysis (Fig S1). The knockout was subjected to a panel of functional analysis, including longevity assay, head swings assay, heat stress assay, and high osmotic stress assay. The results demonstrated that worms’ adaptivity to environment was significantly improved by *CeEngase* knockout (Fig 2).

Recombinant CeENGASE expressed in *E. coli* was prepared for *in vitro* assay. N1H7 was detected and this activity was significantly inhibited by Rabeprazole (Fig 5A). The result was further supported by an *in vivo* test (Fig 5B). Rabeprazole is a proton pump inhibitor used for stomach disease [36–38]. It was reported to have an inhibitory effect on hENGASE [27]. Our study provided a set of new and independent data to support the hypothesis that ENGASE may serve as a new anti-stress and/or anti-aging target.

Both glycan analysis and glycopeptide analysis have being conducted in *C. elegans* since 2003 [39–42]. A preliminary glycomics analysis for N-glycopeptides was conducted for total proteins produced in wild type and knockout worms. The N2H7 glycopeptides were relatively accumulated in *LXR1* worms suggesting N2H7 was a selective substrate for CeENGASE *in vivo* (Table 2), and this was consistent with an *in vitro* result (Fig 5A). N1H0 glycopeptide was not detected in the wild type, indicated that CeENGASE was engaged in a dynamic process.

Basement membrane weas a network existed in multicellular organisms to connect all non-circulation tissues. It was composed of laminin, perlecan, type IV collagen and other specific proteins in *C. elegans* [43]. Early studies had demonstrated that basement membrane was involved in the regulation of cell polarity, survival and tissue organization [44, 45]. Previous study reported a new function of laminin in the regulation of protein toxicity [34]. Our glycomics analysis illustrated that LAM-1, LAM-2, and EPI-1 were relatively more sensitive to CeENGASE regulation, indicating a potential role of CeENGASE in the regulation of structure and functions of basement membrane (Table 1; Fig 6). These observations need to be confirmed by more studies.

