## Supplemental Figure 1 for "Down regulation of *Engase* in *Caenorhabditis elegans* may improve its stresses adaptivity"

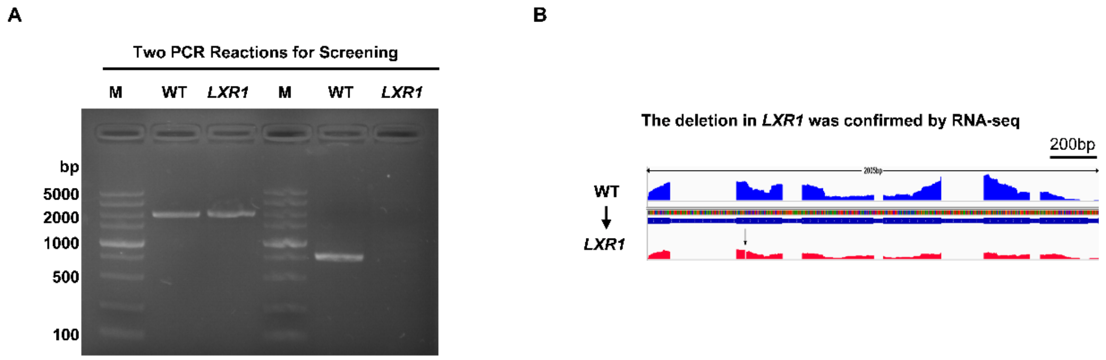


**Figure S1. Construction and validation of a *CeEngase* knockout strain.** (A) The results of two PCR reactions for mutant screening PCR reactions surrounding the targeted mutation site and on deletion site were designed and conducted for sequencing confirmation and quick screening respectively. (B) The deletion in *LXR1* was further confirmed by RNA sequencing analysis. The arrow was the 4bp deletion.
