## Supplemental Table 1 for "Down regulation of *Engase* in *Caenorhabditis elegans* may improve its stresses adaptivity"

**Table S1. Strains used in this study**

| **Strain name** | **Genotype** |
|---|---|
| N2 | wild type |
| LXR1 | *CeEngase(eng-1)*III |
| LXR2 | *PCeEngase::CeEngase::GFP* |
| LXR3 | *PCeEngase::CeEngase(LXR1)::GFP* |

### Table S2. The information for plasmids and primers used in this study

| Construct name | Detailed information | Primers | | Vector | Comments/Reference |
|---|---|---|---|---|---|
| pLXR1 | *CeEnagse [LXR1 (eng-1)]* Cas9-sgRNA | Forward Primer | CACTCGAAGAGCTCTGGAGTgttttagagctagaaatagcaagt | pDD162 | pDD162 was described in (Dickinson et al., 2015). This plasmid was constructed using the protocol of Dickinson 2015. |
| | | Reverse Primer | ACTCCAGAGCTCTTCGAGTGcaagacatctcgcaatagg | | |
| pLXR2 | *PCeEngase::CeEngase::GFP* | Forward Primer 1 (*PCeEngase*) | CGCCAAGCTTtacgatcagctcgatcc | pSM | The *CeEnase (eng-1)* promoter was 2 kb upstream the start codon. |
| | | Reverse Primer 1 (*PCeEngase*) | gctgatcgtaAAGCTTGGCGTAATCATG | | |
| | | Forward Primer 2 (*GFP*) | TGATTCTAATATGAGTAAAGGAGAAGAAC | | Gateway system is published in (Basherudin and Curtis, 2006) |
| | | Reverse Primer 2 (*GFP*) | CTTTACTCATATTAGAATCAATGGAAAAG | | |
| pLXR3 | *PCeEngase::CeEngase(LXR1)::GFP* | Forward Primer | ctcgtatgccacgatatggtggatatttggaagaggAGTCTACGGAAGGAAAGAAATTCG | pSM | A gift from the lab of Kiyoji Nishiwaki. |
| | | Reverse Primer | cctcttccaaatatccaccatatcgtggcatacgagGATTTGACGACCCGAATGTGGTctg | | |
| / | *CeEnagse [LXR1 (eng-1)]* A fast PCR reaction | Forward Primer 1 *(CeEngase)* | ctgctgtcgatgtggcacctctacc | | |
| | | Reverse Primer 1 *(CeEngase)* | gcttaaaatttgaggctgaagctgca | | |
| | | Forward Primer 2 *(CeEnagse [LXR1 (eng-1)] )* | ctgctgtcgatgtggcacctctacc | | |
| | | Reverse Primer 2 (*CeEnagse [LXR1 (eng-1)] )* | AAATATCCACCACGCATATC | | |
| / | *CeEngase (eng-1) RNAi* | Forward Primer | AAGGAATTCACTTCTGGTCC | pPD129.36(L4440) | The full length of *CeEnase (eng-1)* cDNA . |
| | | Reverse Primer | GTGAATAGTAGAAATTTTCG | | |
